# Induction of AmpC-mediated β-lactam resistance requires a single lytic transglycosylase in *Agrobacterium tumefaciens*

**DOI:** 10.1101/2020.09.10.292490

**Authors:** Wanda M. Figueroa-Cuilan, Matthew Howell, Christopher Richards, Amelia Randich, Akhilesh K. Yadav, Felipe Cava, Pamela J.B. Brown

## Abstract

The remarkable ability of *Agrobacterium tumefaciens* to transfer DNA to plant cells has allowed the generation of important transgenic crops. One challenge of *A. tumefaciens*-mediated transformation is eliminating the bacteria after plant transformation to prevent detrimental effects to plants and the release of engineered bacteria to the environment. Here we use a reverse genetics approach to identify genes involved in ampicillin resistance with the goal of utilizing these antibiotic-sensitive strains for plant transformations. We show that treating *A. tumefaciens* C58 with ampicillin led to increased β-lactamase production, a response dependent on the broad-spectrum β-lactamase AmpC and its transcription factor AmpR. Loss of the putative *ampD* orthologue, *atu2113*, led to constitutive production of AmpC-dependent β-lactamase activity and ampicillin resistance. Finally, one cell wall remodeling enzyme, MltB3, was necessary for the AmpC-dependent β-lactamase activity and its loss elicited ampicillin and carbenicillin sensitivity in the *A. tumefaciens* C58 and GV3101 strains. Furthermore, GV3101 *ΔmltB3* transforms plants with comparable efficiency to wildtype but can be cleared with sub-lethal concentrations of ampicillin. The functional characterization of the genes involved in the inducible ampicillin resistance pathway of *A. tumefaciens* constitutes a major step forward in efforts to reduce the intrinsic antibiotic resistance of this bacterium.

**IMPORTANCE:** *Agrobacterium tumefaciens*, a significant biotechnological tool for production of transgenic plant lines, is highly resistant to a wide variety of antibiotics, posing challenges for various applications. One challenge is the efficient elimination of *A. tumefaciens* from transformed plant tissue without using levels of antibiotics that are toxic to the plants. Here, we present the functional characterization of genes involved in β-lactam resistance in *A. tumefaciens.* Knowledge about proteins that promote or inhibit β-lactam resistance will enable the development of strains to improve the efficiency of *Agrobacterium-*mediated plant genetic transformations. Effective removal of *Agrobacterium* from transformed plant tissue has the potential to maximize crop yield and food production, improving the outlook for global food security.

## INTRODUCTION

Rhizobiaceae is a family of bacteria that include soil-dwelling and plant associated bacteria. While some species of this family have the ability to establish symbiotic relationships with plants, others are pathogenic such as the genus *Agrobacterium*. Members of this genus are responsible for a number of diseases, including the cane gall disease (*Agrobacterium rubi*), hairy root disease (*Agrobacterium rhizogenes*), crown gall disease of grapes (*Agrobacterium vitis*), and crown gall disease to flowering plants and woody shrubs (*Agrobacterium tumefaciens*) (1–5). In nature, *A. tumefaciens* causes crown gall by adhering to wounded plants and injecting a section of a bacterial DNA plasmid (Transfer DNA [T-DNA]) that integrates into the plant chromosomes (1–3, 6–11). Expression of genes on the T-DNA segment causes the plant to produce custom energy sources that only *Agrobacterium* can use (9, 10). The increased abundance of energy sources within plant cells leads to their over-proliferation and eventual gall formation (6). Gall formation on plants and trees leads to crop damage, and significant economic losses have been attributed to this issue every year (2, 3).

While the genus *Agrobacterium* exhibits pathogenicity against plants, the natural ability of *Agrobacterium* to transfer DNA to plants has been exploited to produce transgenic plants through genetic engineering (6, 9, 11–13). However, one challenge for *A. tumefaciens*-mediated plant transformations is the elimination of the bacteria from the transformed plant tissue. Elimination of recombinant *A. tumefaciens* from plant tissues is crucial to prevent detrimental effects to plants and to reduce the risk of releasing engineered bacteria into the environment (14–16). β-lactam antibiotics are frequently applied during plant transformations to eliminate *A. tumefaciens* from plant tissues and are preferred over other classes of antibiotics (17–19). Because β-lactams target cell wall synthesis, a process unique to bacteria, and they are less toxic to eukaryotic plant cells than antibiotics that inhibit protein or nucleic acid synthesis (20, 21). However, the natural resistance of *A. tumefaciens* to β-lactams can be only overcome with toxic levels (∼200-1000 mg/L), which has been shown to cause embryogenic tissue necrosis, or to affect plant tissue growth and regeneration rates in a wide variety of plants (17, 18, 22–28). Moreover, depending on the concentration and class of β-lactam, clearing *Agrobacterium* from embryos can take up to 60 days, yet, in some cases, complete elimination of *A. tumefaciens* is not achieved (29). Thus, currently, there is a need for the identification and understanding of regulatory pathways and enzymes involved in β-lactam resistance in *A. tumefaciens.* Functional characterization of bacterial enzymes involved in β-lactam resistance will permit the development of tools that could improve the efficiency of plant genetic transformations, and therefore maximize crop yields and food production.

β-lactam antibiotics target the bacterial cell wall by inhibiting the activity of Penicillin Binding Proteins (PBPs); the enzymes involved in the synthesis of the bacterial peptidoglycan (PG) cell wall (30–38). The bacterial PG cell wall is an essential polymer consisting of alternating *N-*acetylglucosamine (Glc*N*Ac) and *N-*acetylmuramic acid (Mur*N*Ac) sugars crosslinked through peptide bridges (39–44). Because the PG cell wall is a covalently-enclosed polymer, its expansion not only requires cell wall synthesis but also remodeling. Cell wall remodeling is mediated by PG degradation enzymes such as the lytic transglycosylases (LT) (45–48). To allow cell wall expansion, LTs cleave between the Mur*N*Ac and Glc*N*Ac sugar strands resulting in the formation of 1,6-anhydroMurNAc Glc*N*Ac on glycan strands and the liberation of 1,6-anhydromuropeptides (AnhMP) cell wall degradation fragments. The liberated AnhMP fragments are transported to the bacterial cytoplasm for cell wall recycling (42, 49, 50). In the cytoplasm, the recycling of AnhMP fragments keeps the concentration of these products low (51, 52). However, cell wall stressors such as treatment with β-lactam antibiotics or mutations that inhibit the cell wall recycling pathway result in the accumulation of AnhMP cell wall degradation fragments and derepression of β-lactamases (34, 50, 53–56). In bacteria including *Pseudomonas aeruginosa* and *Enterobacter cloaceae*, the AnhMP cell wall degradation fragments are transcriptional activators of inducible β-lactamases, which are enzymes that cleave and inactivate β-lactam antibiotics (57–63).

In the soil environment, many soil microorganisms produce antibiotics to compete for survival, selecting for intrinsic resistance pathways in soil pathogens. For example, the genomes of many soil bacteria contain β-lactamases, such as the cephalosporinase AmpC (34, 64). As a cephalosporinase, AmpC is known to destroy β-lactam antibiotics including monobactams, cephalosporins, and penicillins (34). The AmpC consensus protein sequence consists of a signal sequence for periplasmic transport and a β-lactamase catalytic domain **(Fig. S1A)**. The regulation of AmpC expression varies across bacteria. In *Escherichia coli*, AmpC is a non-inducible β-lactamase that is expressed at low levels and regulated by a promoter and a growth rate-dependent attenuator mechanism (65–67). In contrast, in *P. aeruginosa* and some enterobacteria, AmpC is normally expressed at low levels, but is inducible and can be derepressed during exposure to β-lactams (34, 60, 68). In these cases, AmpC expression is regulated by AmpR, a LysR-type transcriptional regulator found in an operon with AmpC (60). AmpR consists of two domains; a Helix-Turn-Helix DNA-binding domain (DBD) that binds the intergenic region between AmpC and AmpR, and a LysR effector-binding domain (EBD), which contains the regulatory region of AmpR (Fig. 1A**, Fig. S1A**) (57). AmpR is a bifunctional transcriptional regulator that controls both the activation and repression of AmpC (60). The induction mechanism of AmpC by AmpR in response to β-lactams is linked to bacterial cell wall synthesis, remodeling, and recycling (34, 50, 53, 57, 62, 69). Indeed, the AnhMur cell wall degradation fragments released by LTs during cell growth, are AmpR-activating molecules. In contrast, cell wall building blocks such as UDP-Glc*N*Ac Mur*N*Ac pentapeptides bind to AmpR and repress *ampC* transcription (Fig. 1A). A block in bacterial cell wall synthesis after exposure to β-lactams results in accumulation of AnhMur cell wall degradation products in the bacterial cytoplasm, displacement of the AmpR repressor UDP-Glc*N*Ac Mur*N*Ac pentapeptide, activation of AmpR, and transcription of *ampC* (Fig. 1A).

**FIG 1.**
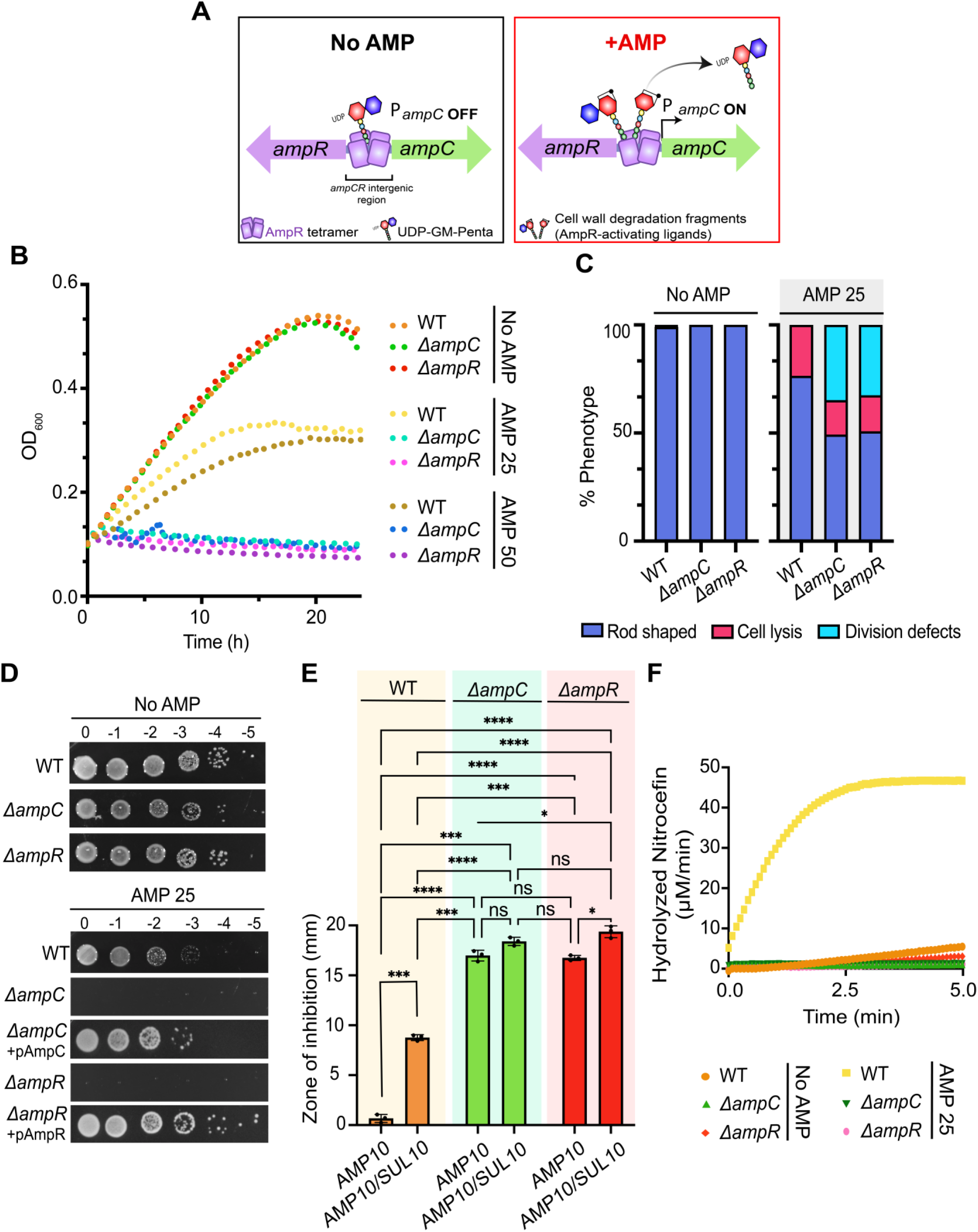
The AmpC-AmpR operon is responsible for induced ampicillin resistance in *Agrobacterium tumefaciens* C58. **(A)** Operon organization and proposed ampicillin resistance mechanism (53). Briefly, in the absence of β-lactams such as ampicillin (AMP), *ampC* expression is repressed by AmpR. AmpR-mediated repression is maintained as long as the AmpR-inactivating ligand, UDP-GM-Pentapeptide, is bound to AmpR (“P*ampC* OFF”). In contrast, the presence of ampicillin (AMP) increases the pools of AmpR-activating ligands, or the cell wall degradation fragments (anhydro modification is depicted by ring), which are known to displace AmpR-inactivating ligands. As a result, the increase of AmpR-activating ligands activates AmpR and *ampC* is transcribed (“P*ampC* ON”). **(B)** Growth of WT *A. tumefaciens, ΔampC*, and *ΔampR* in the absence (No AMP) and presence of ampicillin 25 or 50 µg/ml (AMP 25, AMP 50) for 24 h (n = 1, 2 replicates). **(C)** Quantitative analysis of phase-contrast microscopy of exponentially-growing strains in absence (No AMP) or presence of ampicillin 25 ug/ml (AMP 25). % Phenotype was calculated by counting the number of cells displaying one of the phenotypes indicated (1 cell = 1 phenotype) and dividing by the total number of cells per strain. **(D)** Ampicillin susceptibility assay performed via spotting dilutions. Briefly, indicated strains were cultured overnight (ON) on LB medium, serially diluted, spotted on LB solid medium containing no ampicillin (No AMP) or ampicillin 25 µg/ml (AMP 25), and incubated at 28°C for 36 h before imaging. **(E)** Disc susceptibility was performed on lawns of indicated strains grown on LB plates for 24 h at 28°C (n = 2). AMP 10 = disc containing 10 µg/ml ampicillin, AMP 10/SUL 10 = disc containing 10 µg/ml ampicillin and 10 µg/ml sulbactam, a broad-spectrum β-lactamase inhibitor. Data represent the mean (± standard deviation, SD) of three independent experiments. **** = *P <* 0.0001; *** = *P <* 0.001; ** = *P <* 0.01; * = *P <0.1;* ns = not significant. **(F)** Determination of β-lactamase production performed via a nitrocefin assay using cell lysates. “No AMP” or “AMP 25” indicates cells untreated or treated, respectively, with ampicillin 25 µg/ml for 2 h before the generation of cell lysates. Data shown represents one out of two biological replicates.

The genus *Agrobacterium* is naturally resistant to β-lactams and the molecular basis for this resistance is poorly understood. A study that screened for β-lactamase production in *A. tumefaciens* detected cephalosporinase production (70) and identified one putative cephalosporinase, an AmpC homolog (*atu3077)*, in the *A. tumefaciens* genome. In addition, similar to *Enterobacteriaceae* and *Pseudomonadales, ampC* is found in an operon with *ampR* in most *Agrobacterium* genomes (Fig. 1A). Here, we sought to determine if AmpC is a functional β-lactamase, if AmpC is inducible, and if the natural ampicillin resistance observed in *A. tumefaciens* is dependent on AmpC.

We present a functional characterization of proteins involved in intrinsic ampicillin resistance in *A. tumefaciens.* We find that AmpR is required for AmpC-dependent β-lactamase activity and that loss of the anhydro-amidase AmpD (*atu2113*, misannotated as an AmiD homolog in the genome (71)) leads to increased resistance to ampicillin, a process dependent on AmpC. We suggest that AmpD is required for proper recycling of cell wall degradation products and its loss results in the accumulation of cell wall degradation products and activation of AmpC by AmpR. Furthermore, we find that a single LT, the membrane-bound lytic transglycosylase B3 (MltB3), is necessary for AmpC-dependent β-lactamase activity and its loss leads to ampicillin sensitivity in the *A. tumefaciens* C58 and GV3101 strains. Finally, transformation of *Arabidopsis thaliana* utilizing a *ΔmltB3* GV3101 strain requires significantly lower concentrations of ampicillin while exhibiting similar WT transformation efficiency. This work underscores the significance of understanding the β-lactam resistance pathway of *A. tumefaciens* with the aim of expanding tools for the *A. tumefaciens-*mediated transformations.

## RESULTS AND DISCUSSION

### The AmpC-AmpR operon is responsible for inducible ampicillin resistance in *A. tumefaciens* C58

To begin our characterization, we first assessed the susceptibility of *A. tumefaciens* to different concentrations of ampicillin near the minimum inhibitory concentration (MIC) reported for *A. tumefaciens* on solid and liquid media (72). We found that cells grown in LB with 25 or 50 μg/mL ampicillin (AMP 25 or AMP 50, respectively) for 24 h displayed slow growth in liquid medium in comparison to cells grown in LB without ampicillin (No AMP) (Fig. 1B). To better understand the cause of this growth defect, we performed phase-contrast microscopy of cells treated with AMP 25 for 2 h **(Fig. S1B)**. We found that treatment with AMP 25 causes a significant increase in the median cell length **(Fig. S1C)** and that 23.5% of the cells were undergoing cell lysis (Fig. 1C**, Fig. S1B)**, confirming that the bactericidal effect of AMP 25 on WT *A. tumefaciens* is the cause of the overall decrease in optical density. Similarly, WT cells grown on AMP 25 solid medium for 36 h have a viability defect in comparison to WT cells grown in LB No AMP (Fig. 1D). The increased sensitivity of WT *A. tumefaciens* to ampicillin in the presence of sulbactam, a broad spectrum β-lactamase inhibitor, suggests that β-lactamase production is responsible for the observed ampicillin resistance (Fig. 1E). Finally, to determine if β-lactamase production is induced, we treated WT cells with AMP 25 for 2 h, generated whole cell lysates, and performed nitrocefin assays on total protein content (Fig. 1F). Nitrocefin is a chromogenic substrate related to the cephalosporins that undergoes a color change when is hydrolyzed by β-lactamases (73). After treatment of WT cells with AMP 25 for 2 h, the activity of β-lactamases is readily detected in lysates using nitrocefin assays (Fig. 1F). Together, these results suggest that *A. tumefaciens* C58 β-lactamase production is induced in the presence of β-lactams such as ampicillin. To assess the contributions of putative enzymes involved in ampicillin resistance, we employed a reverse genetics approach (74).

The *ampC* ortholog of *A. tumefaciens* C58 (*atu3077)* is present on the linear chromosome and encodes the only putative inducible β-lactamase in the genome of *A. tumefaciens* C58 (34)*. ampC* is syntenic with *ampR* and the *A. tumefaciens* AmpC and AmpR proteins are 74.7% and 85.9% similar to their respective orthologs from *P. aeruginosa*. To determine the role of AmpC, we deleted *ampC* (*atu3077*) from the *A. tumefaciens* C58 genome. Deletion of *ampC* does not impact cell growth and cell viability (Fig. 1B, 1D) or cell morphology **(Fig. S1B)**, beyond a slight increase in cell length **(Fig. S1C).** To pinpoint the contribution of *ampC* to ampicillin resistance, we assessed the growth dynamics of *ΔampC* cells in the presence of AMP in liquid (Fig. 1B). *ΔampC* cells treated with AMP 25 or AMP 50 for 24 h show a severe growth defect indicating that AmpC contributes to ampicillin resistance. Similarly, deletion of *ampC* results in severe growth viability defect on solid medium containing AMP 25 (Fig 1D). Production of plasmid-encoded AmpC in *ΔampC* restores growth and viability in the presence of AMP 25 (Fig 1D). In addition, *ΔampC* cells treated with AMP 25 exhibit cell division defects (34.8%) and cell lysis (15.8%) (Fig. 1C**, Fig. S1B).** The increased cell length of *ΔampC* treated with AMP 25 is likely due to the observed cell division defects **(Fig. S1B-C**, Fig 1C).

To confirm that ampicillin resistance is mediated by the AmpC β-lactamase, we used the disc diffusion assay to compare resistance levels to ampicillin in the presence and absence of the broad-spectrum β-lactamase inhibitor, sulbactam (Fig. 1E). As expected, *ΔampC* leads to increased sensitivity to ampicillin and the presence of sulbactam does not result in large increases in the zone of growth inhibition (Fig. 1E). Furthermore, monitoring the rates of nitrocefin hydrolysis show that production of β-lactamase is readily detected in WT cells treated with AMP 25 but is undetectable in *ΔampC* following AMP25 treatment (Fig. 1F). Together, these observations suggest that the natural resistance to ampicillin depends on the presence of AmpC, which functions as an inducible β-lactamase.

We hypothesized that if transcription of *ampC* is strictly controlled by AmpR, deletion of *ampR* should mimic deletion of *ampC.* To test this hypothesis, we deleted *ampR* (*atu3078*) from the genome of *A. tumefaciens* C58. Deletion of *ampR* does not impact cell growth (Fig. 1B), morphology **(Fig. S1B)**, cell length **(Fig. S1C)** or cell viability (Fig. 1D) in LB. Low concentrations of ampicillin in either liquid or solid medium are lethal to *ΔampR* and results in similar cell division defects and cell lysis as *ΔampC* (Fig. 1B-C**, Fig. S1B)**. Production of plasmid encoded AmpR restores the viability of *ΔampR* on solid medium with AMP25 (Fig 1D). Like *ΔampC, ΔampR* fails to produce detectable β-lactamase activity when treated with AMP 25 (Fig. 1F). Together, these results suggest that AmpR and AmpC contribute to ampicillin resistance in *A. tumefaciens*. Based on agreement with the general mechanism of characterized AmpR-AmpC pathways, we hypothesize that AmpR is necessary for induction of the AmpC β-lactamase in the presence of ampicillin.

### Loss of AmpD derepresses β-lactamases in *A. tumefaciens* C58

The finding that AmpC and AmpR are necessary for ampicillin resistance in *A. tumefaciens* C58 led us to explore how the pools of different cell wall fragments would alter the AmpC-mediated β-lactamase induction. Similar to exposure to β-lactams, loss of cell wall recycling amidases has been shown to increase the AmpR-activating fragments (cell wall degradation fragments) in the cytoplasm, resulting in the transcriptional derepression of *ampC*and β-lactam resistance (48, 75–78). The genome of *A. tumefaciens* contains one 1,6-anhydro amidase ortholog, *atu2113*, reannotated here as AmpD. The domain organization of AmpD consists of the Amidase_2 (Ami_2) catalytic domain and a PG-binding domain (PBD) that facilitates the interaction with cell wall products **(Fig. S2A)** (45, 48). *A. tumefaciens* AmpD exhibits 64.4% sequence similarity to AmpDh2, one of three broad-spectrum 1,6-anhydro amidase AmpDh paralogs found in *Pseudomonas aeruginosa* (54, 56, 77, 79).

Given that in *A. tumefaciens* ampicillin triggers the AmpC-dependent production of β-lactamases, we hypothesized that if AmpD was an anhydro amidase involved in the recycling of cell wall degradation fragments, its loss should result in increased AmpR-activating fragments in the cytoplasm, β-lactamase induction, and ampicillin resistance. First, we found that *ΔampD* cells exhibit normal cell viability (Fig. 2A), cell growth (Fig. 2B), and morphology **(Fig. S2B-C).** *ΔampD* is highly resistant to ampicillin (Fig. 2A). Indeed, WT cells spotted on AMP 160 are not viable, whereas *ΔampD* spotted on AMP 160 only displays a ∼10-fold decrease in viability compared to untreated cells. In contrast, production of *ampD* from an IPTG-inducible plasmid (+pAmpD) resulted in a 100,000-fold decrease in viability in the presence of AMP 100 (Fig. 2A). In liquid, *ΔampD* cells continue to grow normally, even in the presence of AMP 100 (Fig. 2B) and ampicillin treatment does not trigger obvious morphological changes or cell lysis **(Fig. S2B-C**, Fig. 2C). *ΔampD* produces readily detectable amounts of β-lactamase production in both the presence and absence of ampicillin **(Fig. S2D).** The increased zone of inhibition observed in the presence of ampicillin and sulbactam is consistent with the high level of ampicillin resistance observed in *ΔampD* being mediated by a β-lactamase (Fig. 2D). Together, these results indicate loss of AmpD leads to derepression and increased β-lactamase activity. Our findings are consistent with other bacterial models such as *P. aeruginosa*, where deletion of 1,6-anhydro amidases involved in the recycling of AmpR-activating ligands leads to increased β-lactamase expression (53–56) due to the build-up of activating ligands in the cytoplasm.

**FIG 2.**
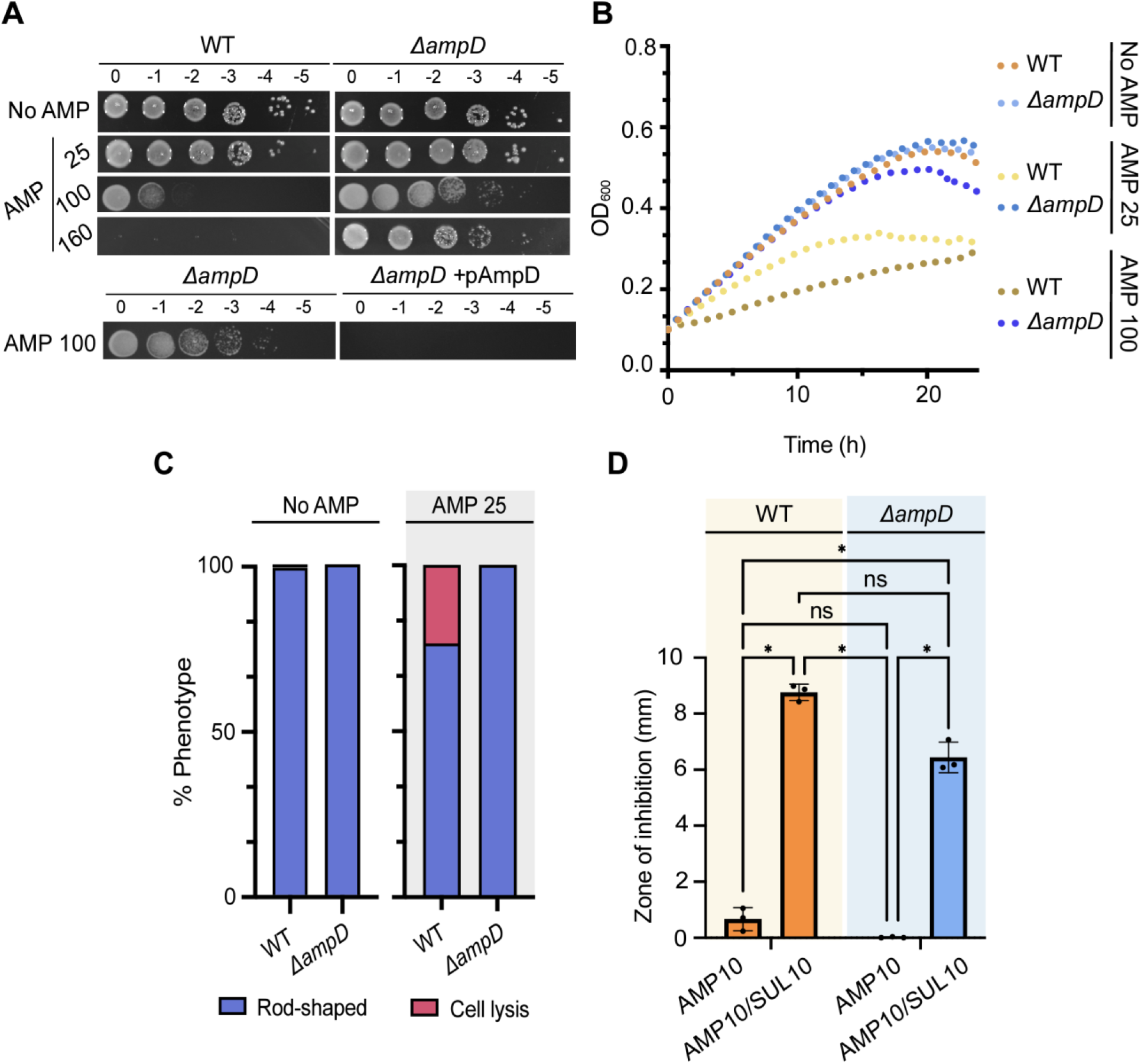
Loss of AmpD results in constitutive β-lactamase activity and elevated ampicillin resistance. **(A)** Ampicillin susceptibility assay performed via spotting dilutions. Briefly, indicated strains were cultured ON at 28°C, serially diluted, spotted on solid medium containing no ampicillin (No AMP) or ampicillin 25, 100, or 160 µg/ml (AMP 25, 100, 160), and incubated at 28°C for ∼40 h before imaging. **(B)** Growth of WT *A. tumefaciens* and *ΔampD* in the absence (No AMP) and presence (AMP) of various concentrations of ampicillin 25, 100 µg/ml (AMP 25, AMP 100) for 24 h (n=1, 2 replicates). **(C)** Quantitative analysis of phase-contrast microscopy of exponentially-growing strains treated with ampicillin 25 µg/ml (AMP 25). % Phenotype was calculated by counting the number of cells displaying one of the indicated phenotypes (1 cell = 1 phenotype) and dividing by the total number of cells per strain. **(D)** Disc susceptibility performed on lawn of indicated strains grown on LB plates for 24 h at 28°C. Blank = no antibiotic disc, AMP 10 = disc containing 10 µg/ml ampicillin, AMP 10/SUL 10 = disc containing 10 µg/ml ampicillin and 10 µg/ml sulbactam, a broad-spectrum β-lactamase inhibitor. Data represent the mean (± standard deviation, SD) of three independent experiments. **** = *P <* 0.0001; *** = *P <* 0.001; ** = *P <* 0.01; * = *P < 0.1;* ns = not significant.

### AmpC is constitutively produced in *ΔampD*

We have shown that AmpC and AmpR are required for ampicillin resistance (Fig. 1), and loss of AmpD leads to elevated β-lactamase activity and ampicillin resistance in *A. tumefaciens* C58 (Fig 2). To confirm that AmpC is the β-lactamase produced by the *ΔampD* strain, we deleted *ampC* or *ampR* in the *ΔampD* background (Fig. 3). In the absence of ampicillin, we found that *ΔampCΔampD* and *ΔampRΔampD* display normal cell viability, growth, and morphology (Fig. 3A-B**, Fig S3A-B).** We found that treatment with AMP 25, on either solid medium or liquid medium, is lethal to *ΔampCΔampD* or *ΔampRΔampD* (Fig. 3A**, Fig. S3A).** Treatment of *ΔampCΔampD* or *ΔampRΔampD* with AMP 25 for 2 h results in lysis of 22.9% and 33.8% of the cells in comparison to untreated cells (No AMP), where lysis is not readily observed (Fig. 3B**, Fig S3B)**. In comparison to *ΔampC* or *ΔampR*, where cell division defects are observed in >20% of the population, very few *ΔampCΔampD* or *ΔampRΔampD* cells exhibit cell division defects when treated with AMP 25 (Fig. 3B). The low incidence of cell division defects observed in *ΔampCΔampD* or *ΔampRΔampD* suggests that the activity of AmpD contributes to the inefficient cell division of *ΔampC* and *ΔampR* cells following ampicillin treatment. Finally, to assess whether the *ΔampD* strain could induce β-lactamase production in the absence of *ampC* or *ampR*, we performed nitrocefin assays. *ΔampCΔampD* and *ΔampRΔampD* fail to produce detectable levels of β-lactamase in the absence or presence of AMP 25 (Fig. 3C). Together, these results suggest that induction of AmpC is the main cause for the elevated resistance to ampicillin observed in *ΔampD*. This data is consistent with the current model for β-lactam resistance in *P. aeruginosa*, where the loss of anhydro amidases leads to accumulation of cell wall degradation products that activate AmpR leading to derepression of AmpC (54–56, 69).

**FIG 3.**
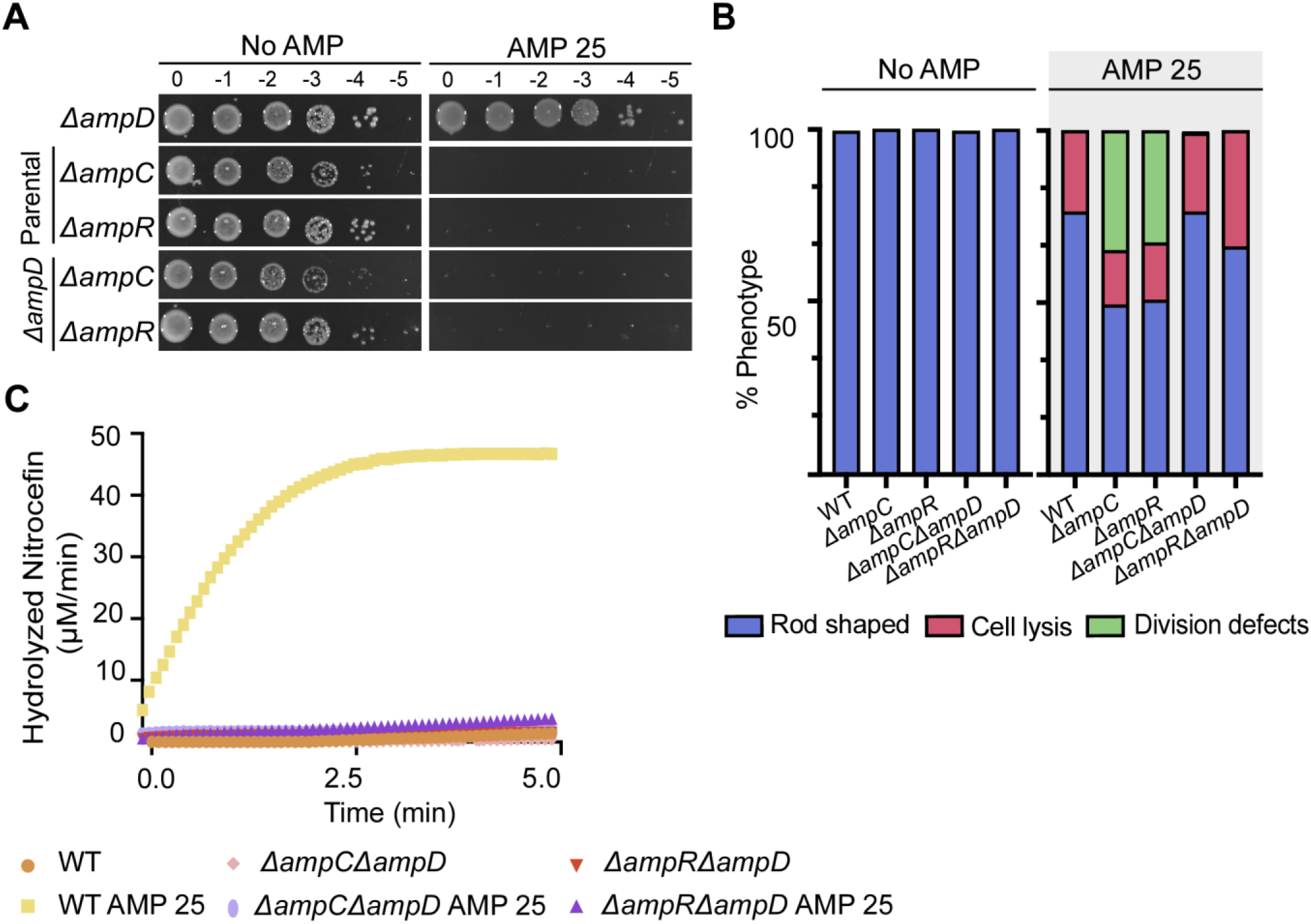
AmpC is the β-lactamase that confers *Δ*ampD elevated ampicillin resistance. **(A)** Ampicillin susceptibility assay was performed via spotting dilutions. Briefly, indicated strains were cultured ON at 28°C, serially diluted, spotted on solid medium containing no ampicillin (No AMP) or ampicillin 25 µg/ml (AMP 25) and incubated at 28°C for ∼40 h before imaging. **(B)** Quantitative analysis of phase-contrast microscopy of exponentially-growing strains untreated (No AMP) or treated with ampicillin 25 µg/ml (AMP 25). % Phenotype was calculated by counting the number of cells displaying one of the indicated phenotypes (1 cell = 1 phenotype). **(C)** Determination of β-lactamase production was performed via a nitrocefin assay using cell lysates. “No AMP” or “AMP 25” indicates cells untreated or treated, respectively, with ampicillin 25 µg/ml for 2 h before the generation of cell lysates. Data shown represents one out of two biological replicates.

### Absence of MltB3 (*Δatu3779*) leads to a failure of AmpC-dependent induction of β-lactamases

Lytic transglycosylases (LTs) are likely to function as the enzymes that generate the AmpR-activating fragments. Different families of LTs have been linked to β-lactam resistance in several bacterial organisms (80, 81). For instance, in *Caulobacter crescentus* deletion of *sdpA*, a soluble LT, led to increased sensitivity to ampicillin (82). In *P. aeruginosa*, loss of several *mltBs* and/or *slt* led to a decrease in the β-lactam MIC, cell viability, and increased outer membrane permeability (83, 84). Thus, we sought to determine if LTs contribute to β-lactam resistance of *A. tumefaciens*. The *A. tumefaciens* genome consists of 8 putative LTs belonging to 3 families: family 1) the soluble lytic transglycosylases (Slt), 2) membrane-bound lytic transglycosylase A (MltA), and 3) membrane-bound lytic transglycosylase B (MltB) **(Fig. S4A)**. We found that single deletions of LTs did not affect cell viability, suggesting a wide redundancy of functions between LTs **(Fig. S4B)**.

Despite the potential for functional redundancy, we found that deletion of a single, family 3, membrane-bound lytic transglycosylase, MltB3 (*atu3779)*, causes ampicillin hypersensitivity **(Fig. S4B)**. Treatment of *ΔmltB3* with AMP 25 for 2 h causes cell lysis defect (28.8%) (Fig. 4A-B) and results in a severe growth defect (Fig. 4C) indicating that MltB3 is required for ampicillin resistance. Production MltB3 from an IPTG-inducible plasmid (+pMltB3) in *ΔmltB3* restores viability in the presence of AMP 25 **(Fig S4B)** confirming that MltB is responsible for this phenotype. Finally, *ΔmltB3* exhibits reduced production of β-lactamase after AMP 25 treatment for 2 hours (Fig. 4D). Together, these results suggest that in *A. tumefaciens*, MltB3 is a specialized enzyme that functions in the AmpR-AmpC β-lactamase pathway. These observations contrast with the *P. aeruginosa* model, in which the β-lactam sensitivity of LT mutants is due to increased outer membrane permeability rather than β-lactamase production (83).

**FIG 4.**
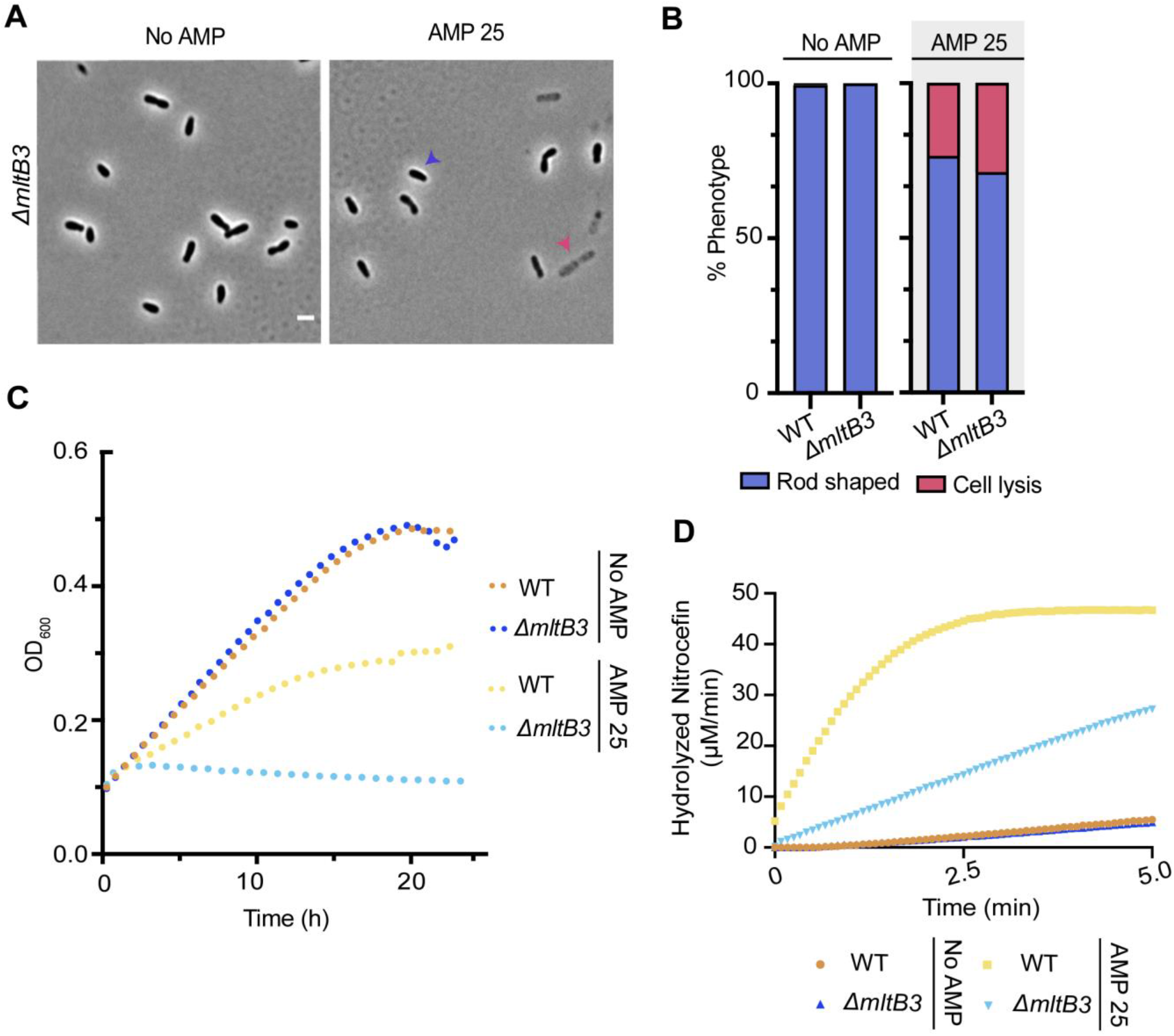
MtlB3 is required for ampicillin resistance in *A. tumefaciens*. **(A)** Phase-contrast microscopy of exponentially-growing strains treated with ampicillin 25 µg/ml (AMP 25) for 2 h. **(B)** Quantitative analysis of phase-contrast microscopy of exponentially-growing strains treated with ampicillin 25 µg/ml (AMP 25) for 2 h. % Phenotype was calculated by counting the number of cells displaying one of the phenotypes indicated (1 cell = 1 phenotype) and diving it by the total number of cells per strain. **(C)** Growth of WT *A. tumefaciens* and *ΔmltB3* in the absence (No AMP) and presence of ampicillin 25 µg/ml (AMP 25) for 24 h (n = 1, 2 replicates). **(D)** Determination of β-lactamase production was performed via a nitrocefin assay using cell lysates. “No AMP” or “AMP 25” indicates cells untreated or treated, respectively, with ampicillin 25 µg/ml for 2 h before the generation of cell lysates. Data shown represents one out of two biological replicates.

### Plant transformation with *Δmltb3* cells requires a low concentration of ampicillin for the elimination of bacteria

Next, we sought to determine if the ampicillin sensitive strains of *A. tumefaciens* constructed in this work are competent for plant transformation. While the *ΔampC* strain appears to be the ideal mutant for these studies, we took into consideration the impact of the mutation on the overall fitness of our ampicillin sensitive strains. *ΔampC* and *ΔampR* growth dynamics are very similar to WT in liquid; however, these strains exhibit a 10-fold viability defect on solid medium. In contrast, *ΔmltB3* cells growth dynamics mimic that to WT and lyse quicky in the presence of low concentrations of ampicillin. Thus, we reasoned that the ΔmltB3 allele would enable us to test the transformation efficiency of an otherwise fit but ampicillin sensitive *A. tumefaciens* strain. To this end, we deleted *mltB3* in *A. tumefaciens* GV3103 and find that this mutation causes susceptibility to AMP 25 and carbenicillin 15 (CARB 15) (Fig. 5A). To confirm that the absence of MltB3 (*ΔmltB3*) prevented the induction of β-lactamase production in the GV3101 strain after ampicillin treatment, we performed nitrocefin assays using lysates of cells treated with AMP 25 for 2 h. While β-lactamase activity is readily detected from WT GV3101-treated cells with AMP 25, the *ΔmltB3* mutant produces relatively low levels of inducible β-lactamase (Fig. 5B). Together, we conclude that MltB3 is the major LT in *A. tumefaciens* C58 and GV3101 that contributes to *A. tumefaciens* natural resistance to ampicillin and other β-lactam antibiotics.

**FIG 5.**
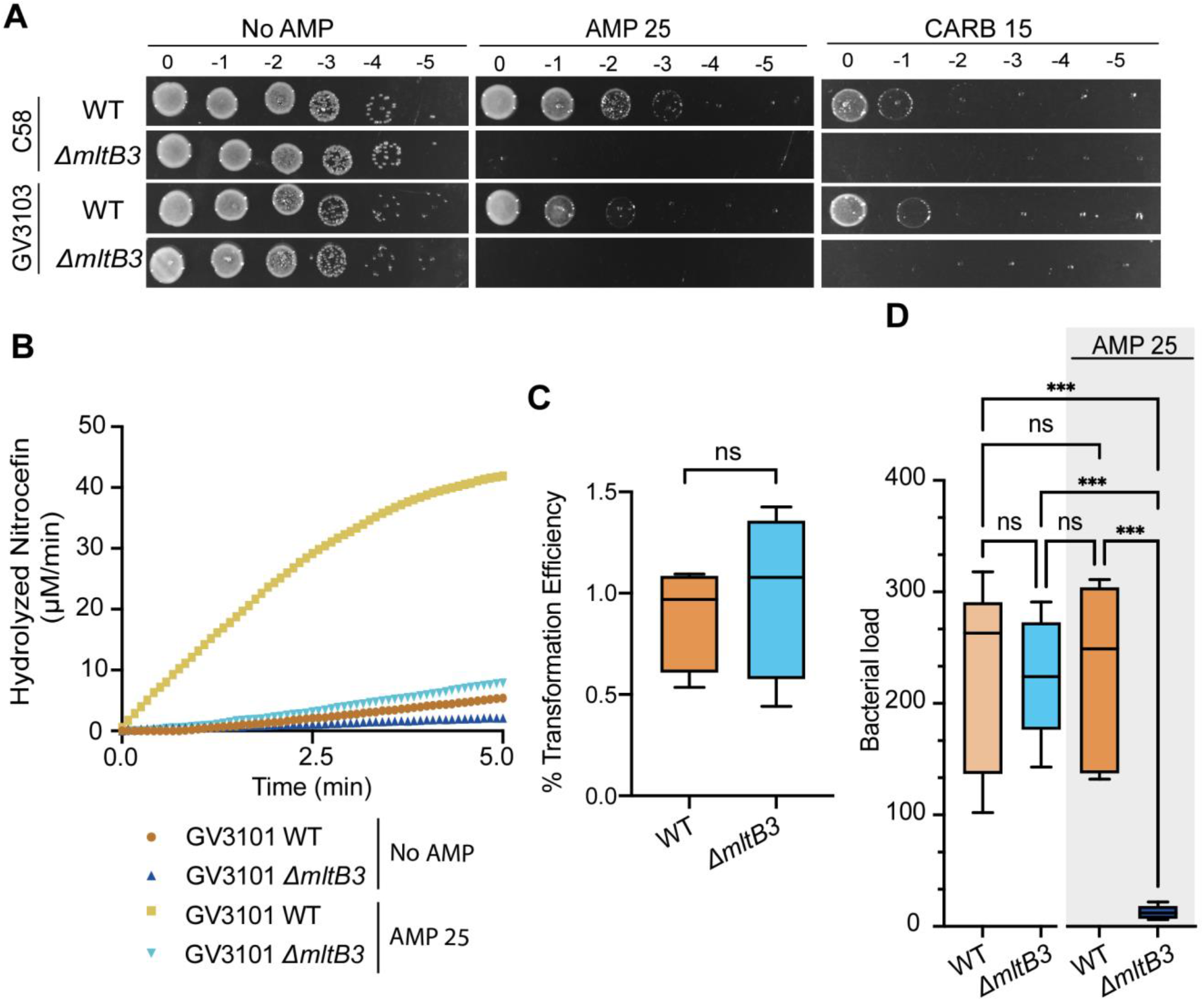
*A. tumefaciens* GV3103 *ΔmltB3* can be used to transform plants efficiently and bacteria can be removed using low concentrations of ampicillin. **(A)** Ampicillin susceptibility assay was performed via spotting dilutions. Briefly, indicated strains were cultured ON at 28°C, serially diluted, spotted on solid medium containing no ampicillin (No AMP), ampicillin 25 µg/ml (AMP 25), or carbenicillin 15 µg/ml (CARB 15), and incubated at 28°C for ∼40 h before imaging. **(B)** Determination of β-lactamase production was performed via a nitrocefin assay using cell lysates. Data shown represents one out of two biological replicates. “No AMP” or “AMP 25” indicates cells untreated or treated, respectively, with ampicillin 25 µg/ml for 2 h before the generation of cell lysates. **(C)** Transformation efficiency and (D) bacterial loads of seeds transformed with WT *A. tumefaciens* GV3101 and GV3101 *ΔmltB3* using the floral dip assay technique (85).

Next, we confirmed that *ΔmltB3 A. tumefaciens* GV3101 effectively transforms *Arabidopsis thaliana* using the standard floral dip technique (85) with an efficiency comparable to WT (Fig. 5C). Next, we asked whether elimination of the bacteria could be performed using low concentrations of ampicillin (AMP 25). Seeds transformed with WT *A. tumefaciens* GV3101 contain similar bacterial loads compared to seeds transformed with *Δmltb3 A. tumefaciens* GV3101 when plated on solid media without ampicillin (Fig. 5D). When plated on media containing AMP 25, seeds transformed with *Δmltb3* exhibit a significant drop in bacterial load. In contrast, this low level of ampicillin does not reduce the bacterial load of seeds transformed with WT GV3101 (Fig. 5D). These results demonstrate that the *Δmltb3* GV3101 strain is useful for the transformation of *Arabidopsis thaliana* and elimination of the bacteria can be performed using lower concentrations of ampicillin. While ampicillin or carbenicillin are occasionally used for clearing *Agrobacterium* after transformation, many labs routinely use expensive antibiotics such as the proprietary blends of Timentin and Augmentin (86, 87). Our work highlights the ability to clear *ΔmltB* cells using ampicillin, a cost-effective and readily available antibiotic. Furthermore, the increased sensitivity of this strain to carbenicillin suggests that bacterial clearance following plant transformation can likely be achieved using other β-lactam antibiotics. Overall, these data show the potential impact of improved understanding the cell biology of *A. tumefaciens* to improve genetic engineering approaches.

## CONCLUDING REMARKS AND FUTURE PERSPECTIVES

The natural ability of *A. tumefaciens* to transform plants has allowed the production of transgenic crops of incredible economic importance for the past four decades. One challenge of the *A. tumefaciens*-mediated plant transformation is natural resistance of *A. tumefaciens* to antibiotics, which requires toxic concentrations of antibiotics to eliminate *A. tumefaciens* from transformed tissues. Here we show that *A. tumefaciens* induces β-lactamase activity in response to ampicillin exposure. Indeed, induction of β-lactamase activity upon exposure to ampicillin is dependent on the β-lactamase AmpC and the transcription factor AmpR. Moreover, we found that deletion of a single LT, MltB3, sensitizes *A. tumefaciens* to the β-lactams.

We propose that during *A. tumefaciens* growth and remodeling, there is a delicate balance between the synthesis and degradation of the bacterial cell wall. PBPs insert precursor cytoplasmic monomers into the growing cell wall polymer (Fig 6**, steps 1-2**). During remodeling, cell wall hydrolytic enzymes such as such as endopeptidases and LTs, including MltB3, liberate cell wall degradation products, which are transported back into the cytoplasm of *A. tumefaciens* for their recycling (Fig 6**, steps 3-4**). In the cytoplasm, hydrolytic enzymes including L,D-carboxypeptidases (LD-CPases), amidases such as AmpD, and glycosidases limit the pool of cell wall fragments by allowing their recycling (Fig 6**, steps 5-8).** When cell wall degradation fragments accumulate during β-lactam treatment, the AmpR-dependent production of the AmpC β-lactamase is increased (Fig 6**, step 9)**. Overall, the identification and contributions of genes conferring ampicillin resistance in *A. tumefaciens* will be beneficial for improving the design of *A. tumefaciens-*mediated genetic engineering.

**FIG 6.**
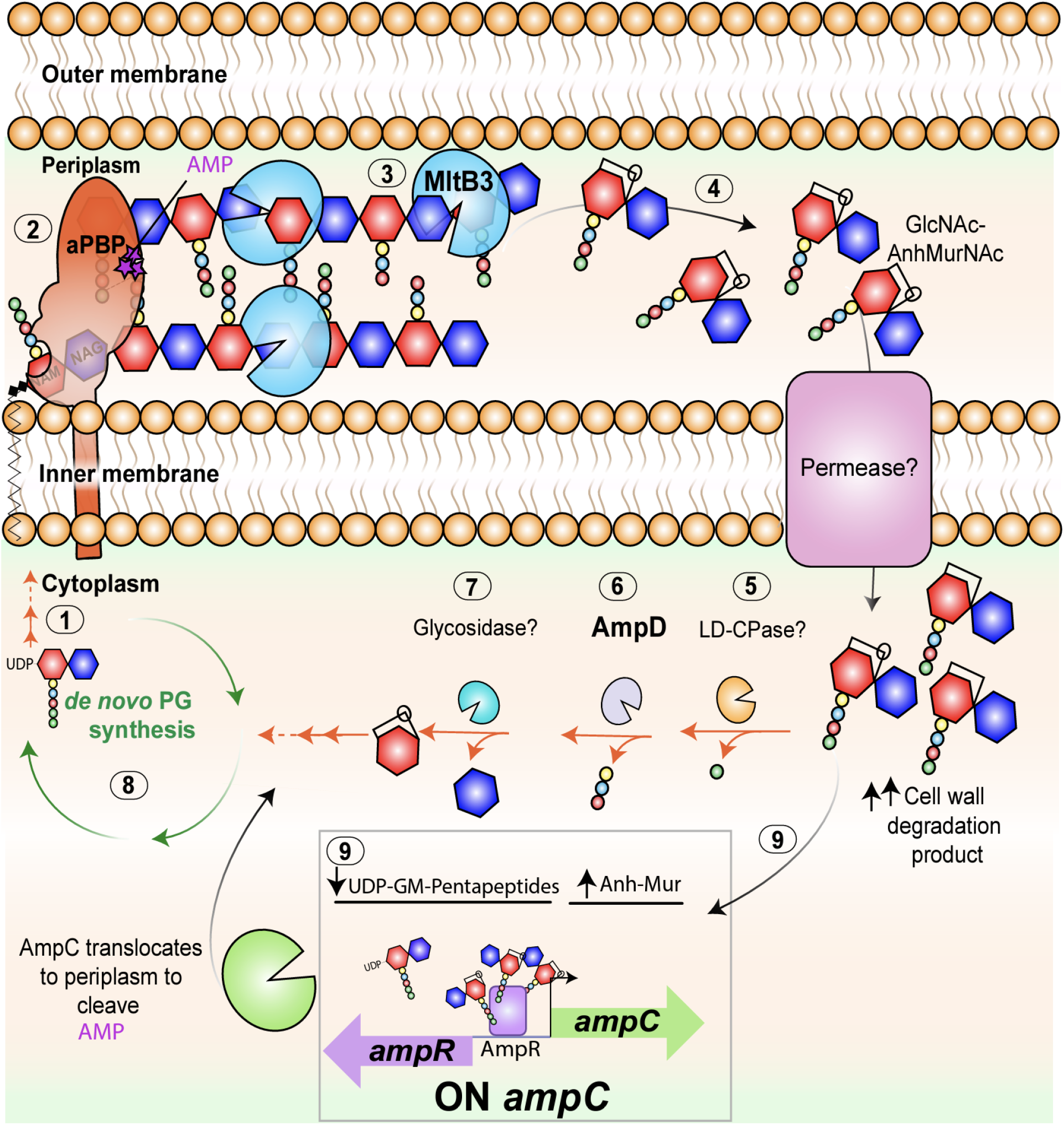
Working model for *A. tumefaciens* ampicillin resistance. Bifunctional PBPs extend the cell wall through the transglycosylation (linking of carbohydrates) and transpeptidation reactions (linking of peptide stems). β-lactams such as ampicillin (purple stars) target the transpeptidase domain of Penicillin-Binding Proteins (PBPs) (**Fig S6**, Step 1-2), leading to a block in bacterial cell growth. In *A. tumefaciens*, treatment with ampicillin leads to induction of β-lactamases. Inactivation of the lytic transglycosylase MltB3 results inhibition of β-lactamase derepression and lysis suggesting that MltB3 is likely required for the generation of cell wall degradation products (**Fig S6**, Step 3). Similar to treatment with β-lactams, where a block in cell growth leads to an increase in cell wall degradation products (**Fig S6**, Step 5), inactivation of anhydro amidases such as AmpD increases the poll of this molecules leading to β-lactam resistance. In *A. tumefaciens* inactivation of AmpD leads to derepression of β-lactamases and ampicillin resistance (**Fig S6**, Step 9). Both AmpC, an inducible β-lactamase that is under the transcriptional control of AmpR, and AmpR seem to be responsible for the derepression observed in *ΔampD.* Thus, our working model suggests that upon ampicillin exposure, a block in growth leads to increased activity of MltB3. An increase in cell wall degradation products leads to induction of AmpC expression by AmpR and the presumed translocation of AmpC to the periplasm resulting in ampicillin resistance.

## MATERIALS AND METHODS

### Bacterial strains and growth conditions

Unless otherwise indicated, *Agrobacterium tumefaciens* GV3101, C58, and derived strains were grown in LB medium (10 g/l tryptone, 10 g/l yeast extract and 5 g/l NaCl) at 28°C with shaking and the antibiotic kanamycin was added at a concentration of 100 μg/ml to maintain plasmids in complementing strains. For determining bacterial loads after transformation, *Agrobacterium tumefaciens* GV3101 was grown in minimal medium ATGN (88). *E. coli* DH5α and S17.1 were routinely cultivated in LB medium at 37°C with shaking and the antibiotic kanamycin was added at a concentration of 50 μg/ml and 25 μg/ml to maintain plasmids in DH5α and S17-1 strains, respectively.

### Construction of plasmids and strains

All strains and plasmids used are listed in **Table 1**. Synthesized DNA primers are listed in **Table 2**.

Gene deletions were achieved by allelic exchange and vectors were constructed as previously described (89). Briefly, 500 base pairs (bp) fragments upstream and downstream of the gene of interest were amplified from purified C58 genomic DNA using primer pairs P1/P2; which amplify 500 bp upstream of the gene of interest, and P3/P4; which amplify the 500 bp downstream of the gene of interest. Overlapping PCR was used to merge and amplify the amplicons generated by P1/P2 and P3/P4, using primer pair P1/P4. The 1,000 bp amplicon was digested and ligated into the deletion plasmid, pNTPS139. The deletion plasmids were introduced into *A. tumefaciens* by mating using an *E. coli* S17.1 conjugation strain to create kanamycin (KAN) resistant, sucrose sensitive primary exconjugants. Deletion strains were constructed as described previously (74). Briefly, primary exconjugants were grown overnight at 28°C in ATGN with no selection and plated in ATGN KAN 300 for 48 h at 28°C. Colonies were screened by patching for KAN resistance and sucrose sensitivity. Colony PCR using primers P5/P4 were used to confirm that recombination took place and at the region of interest. Next, positive colonies are grown in ATGN at 28°C overnight and plated in ATSN 5% sucrose. Secondary recombinants were screened by patching for sucrose resistance and KAN sensitivity. Colony PCR with primers P5/P6 for the respective gene target was used to confirm deletion. PCR products from P5/P6 primer sets were sequenced to further confirm deletions.

For the construction of replicating plasmids, the amplicons and pSRKKM-Plac-*sfgfp*, were digested overnight and ligated overnight at 4°C using NEB T4 DNA ligase and transformed into *E. coli* DH5α. Plasmids were sequenced to verify content and were introduced into *A. tumefaciens* by mating using an *E. coli* S17.1 harboring the appropriate plasmid.

### Cell viability spot assays

The cell viability assay was performed as described (90). For cell viability spot assays, cultures were grown overnight and diluted to an OD_600_ of 0.05 and serially diluted in LB up to 10^-6^. 4 µl of each dilution was spotted onto LB plates and incubated at 28°C for 36-40 h before imaging. To determine ampicillin antibiotic resistance, *A. tumefaciens* cultures were grown overnight and diluted to an optical density at 600 nm (OD_600_) of 0.05 and serially diluted and spotted onto LB plates containing AMP 25, AMP 100, or AMP 160. IPTG-inducible complementing strains were grown overnight in the absence of IPTG and diluted to an OD_600_ of 0.05 and serially diluted and spotted onto LB plates containing 1µM IPTG, Kan 150 µg/ml and AMP 25 or AMP 100.

### Phase-contrast microscopy

For phase-contrast microscopy, 0.8 µl of exponentially-grown cultures were spotted onto a 1.25% agarose pad as previously described (90) using a Nikon Eclipse Ti inverted microscope and imaged using a Nikon Plan 60X oil Ph3 objective. Cell length analysis was performed using the MicrobeJ plug-in for Fiji (91). A One-Way analysis of variance (ANOVA) Kruskal-Wallis test with Dunn’s post-test was used to compare the indicated strains. Images were prepared using Adobe Photoshop, Adobe Illustrator, and Prism.

### Growth curves

For growth curves, exponentially growing cultures were diluted to an OD_600_ = 0.2 and 100 µl of dilute culture was added to wells of a 96-well plate. OD_600_ reading were recording using a plate reader at 28°C with shaking every 5-10 min. When indicated, ampicillin was added to a final concentration of AMP 25 or AMP 100. Plots of OD_600_ data represent two technical replicates for each culture measured every 5 minutes for 24-48 h.

### Disc diffusion assay

The disc-diffusion assay was used to determine the resistance of *Agrobacterium* to various antibiotics. Cells were grown on LB agar plates. Sterile discs (6.5 mm in diameter) were placed on the surface of LB agar plates seeded with overnight culture of indicated strains. We used three different discs: Blank (sterile disc containing no antibiotic), AMP 10 (ampicillin 10µg/ml), and AMP 10/SUL 10 (ampicillin 10µg/ml sulbactam 10µg/ml). The LB agar assay plates used for testing *A. tumefaciens* susceptibility were incubated at 28°C for 24-36 h. The assessment of antibacterial activity was based on the measurement of diameter of the zone of inhibition formed around the disc minus the size of the disc. Three independent trials were conducted for each concentration of each antibiotic. A two-way analysis of variance (ANOVA) was used to compare the means.

### Nitrocefin Assay

Nitrocefin is a chromogenic substrate for measuring β-lactamase activity. Nitrocefin has an absorbance maximum of 390 nm. Upon hydrolysis of the β-lactam ring by a β-lactamase, the absorbance shifts from 390 nm to 486 nm. By monitoring absorbance at 486 nm over time and using Beer’s law (A=εlc), we directly measured the rate of β-lactamase hydrolytic activity. In two separate experiments, indicated strains were grown in LB until desired optical densities (OD600) of 0.6 were reached. The cells were then pelleted by centrifugation at 25,900 x g (rotor: Fiberlite F14-6×250y) for 5 minutes and the supernatant collected and washed three times with PBS. Cell lysates were generated by adding the cell lysis buffer BugBuster and sonicating. No lysozyme or protease inhibitors were added. Cell lysates were normalized based on total protein content (7.5 µg/ml) and volume before incubation in 100 μM nitrocefin solution in a 200 μl reaction volume at room temperature in 20 mM HEPES, 300 mM NaCl, pH 7.5. BugBuster lysis buffer was used as a blank. *ΔampD* lysates were normalized based on total protein content (7.5 µg/ml) and subsequently diluted to 1:5 as the rate of nitrocefin hydrolysis was significantly faster than the controls (WT, WT AMP 25). Absorbance was immediately measured at 486 nm in 5 second intervals for 300 seconds (5 minutes). The change in absorbance (A) was converted to change in concentration (c) of hydrolyzed product by using Beer’s law (A=εlc) where the molar extinction coefficient (ε) = 20500 M-1 cm-1 and path length (l) = 1 cm. Data plots of one experiment appears in the main figures and the other in the supplementary figures respectively.

### Construction of Agrobacterium strain for plant transformation experiments

*mltB3* was deleted from the genome of WT *Agrobacterium tumefaciens* GV3101 using the same plasmid and technique used to delete the gene in the strain C58. The resulting strain, GV3101 *ΔmltB3*, was next transformed by introduction of the empty binary vector, pUBQ10-GW (92) via electroporation. This plasmid confers KAN resistance to the *Agrobacterium* strain, and also contains T-DNA repeat blocks which allow the transfer of BASTA resistance to transformed into *Arabidopsis thaliana*. pUBQ10-GW was also electroporated into wildtype GV3101 as a control.

### *Arabidopsis thaliana* transformation efficiency

Plants with a bolt height between 2 and 7 centimeters were transformed via the floral dip method (85). 5 plants were transformed on separate days from independent colonies for each C58 and GV3101. Plants were grown in Promix BX (Premier Tech Horticulture) at 23°C, 16 h light/8 h dark, 100–150 μE m^-2^ s^-1^, and 50–70% humidity until seeds were fully developed. Seeds were collected from fully matured plants and store at 4°C at low humidity for one week. Seeds were then surface sterilized in 15% bleach containing 0.1% Triton X-100 with gentle rocking for 5 minutes. Seeds were then washed 3 times using sterile water. Seeds were then imbibed in sterile water at 4°C for 3 days. Seeds were then sown into soil using conditions listed above. Once seeds developed true leaves BASTA (glufosinate ammonium) was applied via spray at a concentration of 10mg/L. After 4 days BASTA was re-applied ensuring only transformed plants survived.

### Determining bacterial load of transformed *Arabidopsis thaliana* seeds

Seeds were collected from fully matured transformed plants and stored at 4°C for 1 week with low humidity. Seeds were then surface sterilized in 15% bleach containing 0.1% Triton X-100 with gentle rocking for 5 minutes. Seeds were then washed 3 times using sterile water. 10mg of seeds for each experimental condition were ground with a sterile mortar and pestle. Ground seeds were then suspended in 1mL of sterile water, and serially diluted. 200µl of 10^-1^ dilution was plated on minimal medium ATGN No AMP and AMP 25. Plates were then incubated at 28°C before colonies were counted.

## ACKNOWLEDGEMENTS

We thank the Brown lab for helpful discussions. Research in the Brown lab on *A. tumefaciens* cell growth is supported by the National Science Foundation (IOS1557806). WFC was supported by a Gus T. Ridgel Fellowship.

## AUTHOR CONTRIBUTIONS

WFC and PJBB conceptualized this work; WFC and CR developed methods; WFC, MH, CR, AMR and AKY performed the experiments and completed data analysis; WFC, AMR, and PJB wrote the manuscript; WFC, AMR, AKY, FC, and PJBB edited the manuscript; PJBB acquired funding for this work.

