## Supplemental Material for "Induction of AmpC-mediated β-lactam resistance requires a single lytic transglycosylase in *Agrobacterium tumefaciens*"

**Running title:** AmpC-mediated b-lactam resistance of *A. tumefaciens*

**[#]Corresponding author:**

(573) 884-0384

**Present addresses:**
*Department of Biological Chemistry, Johns Hopkins University School of Medicine, Baltimore, Maryland 21205
[&]Department of Biology, Westminster College, Fulton, Missouri 65251
[$]Analytical Chemistry Division, CSIR-Indian Institute of Toxicology Research, Lucknow-226001, India
[^]Department of Biology, The University of Scranton, Scranton, Pennsylvania,18510

**Contents**
Supplemental Tables 1-2
Supplemental Figures and Legends 1 – 4

### SUPPLEMENTARY TABLES

**Table S1.** Strains and plasmids used in this study.

| Strain or plasmid | Relevant characteristics | Reference/Source |
|---|---|---|
| ***Source Plasmids*** | | |
| pSRKKm-Plac-sf*gfp* | pSRKKm vector containing lacI$^q$ and *lac* promoter with sfGFP | (90) |
| pNTPS139 | Km$^r$; Suicide vector containing *oriT* and *sacB* | D. Alley |
| ***Plasmids*** | | |
| *Deletion plasmids* | | |
| pNTPS139 *Δatu3077* | Km$^r$ Suc$^s$; deletion plasmid for *atu3077* | This study |
| pNTPS139 *Δatu3078* | Km$^r$ Suc$^s$; deletion plasmid for *atu3078* | This study |
| pNTPS139 *Δatu0092* | Km$^r$ Suc$^s$; deletion plasmid for a*tu0092* | This study |
| pNTPS139 *Δatu2122* | Km$^r$ Suc$^s$; deletion plasmid for a*tu2122* | This study |
| pNTPS139 *Δatu3779* | Km$^r$ Suc$^s$; deletion plasmid for a*tu3779* | This study |
| pNTPS139 Δ*atu2113* | Km$^r$ Suc$^s$; deletion plasmid for *ampD* | This study |
| *Replicating plasmids* | | |
| Plac-*Atu3077* | Km$^r$; pSRKKm vector containing lacIq and *lac* promoter for the production of AmpC | This study |
| Plac-*Atu3078* | Km$^r$; pSRKKm vector containing lacIq and *lac* promoter for the production of AmpR | This study |
| Plac-*Atu2113* | Km$^r$; pSRKKm vector containing lacIq and *lac* promoter for the production of AmpD | This study |
| Plac-*Atu3779* | Km$^r$; pSRKKm vector containing lacIq and *lac* promoter for the production of MltB3 | This study |

| | | |
|---|---|---|
| ***E. coli strains*** | | |
| DH5α | Cloning strain | Life Technologies |
| S17.1 | $Sm^r$; RP4-2, Tc::Mu,Km-Tn7, for plasmid mobilization | (92) |
| ***A. tumefaciens strains*** | | |
| C58 | Nopaline type strain; pTiC58; pAtC58 | |
| GV3101 | C58-derived; pTiC58DT-DNA; strain for *Agrobacterium* mediated transformation of dicots | John Walker Lab |
| FC2452 | Deletion strain for *ΔmltA (atu0009)* in C58 | This study |
| FC2444 | Deletion strain for *ΔmltB1 (atu0092)* in C58 | This study |
| FC2465 | Deletion strain for *ΔmltB2 (atu2122)* in C58 | This study |
| FC2487 | Deletion strain for *ΔmltB3 (atu3779)* in C58 | This study |
| FC2446 | Deletion strain for *Δslt1 (atu1022)* in C58 | This study |
| FC2448 | Deletion strain for *Δslt2 (atu2112)* in C58 | This study |
| FC2450 | Deletion strain for *Δslt3 (atu2117)* in C58 | This study |
| C58 *ΔtetRA*::a-*att*Tn7 | Replacement of the *ΔtetRA* locus with an artificial *att*Tn7 site | (90) |
| C58 *ΔtetRA*:: a-*att*Tn7 *Δatu3077* | Deletion strain for *ΔampC* | This study |
| C58 *ΔtetRA*:: a-*att*Tn7 *Δatu3077* pSRKKm-Plac-*Atu3077* | $Km^r$; Deletion of *ΔampC in* C58*ΔtetRA::*a-attTn7 carrying complementing plasmid | This study |
| C58 *ΔtetRA*:: a-*att*Tn7 *Δatu3078* | Deletion strain for *ΔampR* | This study |
| C58 *ΔtetRA*:: a-*att*Tn7 *Δatu3078* Plac-*Atu3078* | $Km^r$; Deletion of *ΔampR in* C58*ΔtetRA::*a-attTn7 carrying complementing plasmid | This study |

| | | |
|---|---|---|
| C58 Δ*tetRA*::a-*att*Tn7 *ΔampD* | Deletion strain for *ΔampD* | This study |
| C58 Δ*tetRA*::a-*att*Tn7 *ΔampD* Plac-*ΔampD* | Km$^r$; Deletion of *ΔampD in* C58*ΔtetRA::*a-attTn7 carrying complementing plasmid | This study |
| C58 Δ*tetRA*::a-*att*Tn7 *ΔampD ΔampC* | Deletion strain for *ΔampDΔampC* | This study |
| C58 Δ*tetRA*::a-*att*Tn7 *ΔampD ΔampR* | Deletion strain for *ΔampDΔampR* | This study |
| C58 GV3101 *Δmltb3* | Deletion strain for *ΔmltB3* | This study |
| C58 GV3101 *Δmltb3* Plac-*ΔmltB3* | Km$^r$; Deletion of *ΔmltB3 in* C58*ΔtetRA::*a-attTn7 carrying complementing plasmid | This study |

**Table S2**. Synthesized DNA primers used in this study.

| Synthesized DNA | Sequence |
|---|---|
| *atu3077* P1 | 5’-GCGGCGACTAGTAAACGGATGCCGCTTTTGAAATGC-3’ |
| *atu3077* P2 | 5’-AAGCTTGGTACCGAATTCGCGATTAAATTTCATCTTTCGTGT-3' |
| *atu3077* P3 | 5’-GAATTCGGTACCAAGCTTGCGCTCGAAAAGGCGCAATAA-3' |
| *atu3077* P4 | 5’-GTCGTCGGATCCAGATAACTCGGCACACGCCCA-3' |
| *atu3077* P5 | 5’-CTGCGCCGCCGGTGAAACGCCCGC-3' |
| *atu3077* P6 | 5’-TGTGCCGGAGGCGCTTGCGATCGC-3' |
| *atu3078* P1 | 5’-GTCGTCACTAGTGGGTTTTCCCTTCATGGACGG-3' |
| *atu3078* P2 | 5’-AAGCTTGGTACCGAATTCAAATTGCCGAACCATTCAAGACCT-3' |
| *atu3078* P3 | 5’-GAATTCGGTACCAAGCTTGAGACCATCGGAACGGCGTGA-3' |
| *atu3078* P4 | 5’-GTCGTCGGATCCCCTGTTTGATGCTTTTTATCGCGC-3' |
| *atu3078* P5 | 5’-CCAAGCGCCGGGAAAAGCGTGTCC-3' |
| *atu3078* P6 | 5’-GGCATCGTCTGCGTGGTGTTCGTC-3' |
| *atu2113* P1 | 5’-GTCGTCACTAGTCAGATTTTCGATTTCCCGGACGAA-3’ |
| *atu2113* P2 | 5’-AAGCTTGGTACCGAATTCCGAACATTCTTTCATGCGACG-3’ |

| | |
|---|---|
| *atu2113* P3 | 5’-GAATTCGGTACCAAGCTTCTGCCGAGATTTTCGGCCGCCTGA-3’ |
| *atu2113* P4 | 5’-GTCGTCGGTACCCCACCGAAACCACGGCATGCGCCA-3’ |
| *atu2113* P5 | 5’-GCGCAATCGGCGACGCGG-3' |
| *atu2113* P6 | 5’-GTCGTCTGCTGCACACTTCGCCGC-3’ |
| *atu0009* P1 | 5’-AAAAAGCTTAACGCATCTTCTAGCCTTGCG-3’ |
| *atu0009* P2 | 5’-TATTCATTGCTCGGATTCGG-3’ |
| *atu0009* P3 | 5’-GCAATGAATATATCGGCGATGAAAGGC-3’ |
| *atu0009* P4 | 5’-AAAGGATCCGAAAGAACAATTCCTCCGC -3’ |
| *atu0009* P5 | 5’-TTTCGGCGACCTATGACAAGGACGG-3’ |
| *atu0009* P6 | 5’-TTACCAGTTTGCGGAACGCTGGG-3’ |
| *atu0092* P1 | 5’-AAACTGCAGATTCTTGCCCTGATGCCCATTGTCGC-3’ |
| *atu0092* P2 | 5’-AGACCGAATATCGTCTTTTAATGCTGGTCGG-3’ |
| *atu0092* P3 | 5’-ATTCGGTCTCCTCTTGGATGG-3’ |
| *atu0092* P4 | 5’-AAAGGATCCAAGTCGAGATCGACTGAGCCC-3’ |
| *atu0092* P5 | 5’-AATATGTCCGCCACAACCATCGTCGC-3’ |
| *atu0092* P6 | 5’-ATTACATCACAGACCGCCTCTCCG-3’ |
| *atu2122* P1 | 5’-AAAGAATTCAATCATCAGGGTTCCAATGCGG-3’ |

| | |
|---|---|
| *atu2122* P2 | 5’-TAGTGCGATTTTCCTCGATAGGTTGTTGGC-3’ |
| *atu2122* P3 | 5’-ATCGCACTACCGGGCCTTTAATCTATCGG-3’ |
| *atu2122* P4 | 5’-AAAAAGCTTATAATGACGTCTTTGAACGC-3’ |
| *atu2122* P5 | 5’-TATACCGCAACCGGCGTCGTACCCG-3’ |
| *atu2122* P6 | 5’-TTTCTGTGATGCGGTGCAGCACGG-3’ |
| *atu3779* P1 | 5’-AAACTGCAGTAGAAATTCGACGGCGCCG-3’ |
| *atu3779* P2 | 5’-TTTCGATTGCGAAAACGCATCGGGCG-3’ |
| *atu3779* P3 | 5’-GCAATCGAAATAATGTGCCGGCGAATTCGG-3’ |
| *atu3779* P4 | 5’-AAAGGATCCTTGGCCGTTCATGTCGTAGCC-3’ |
| *atu3779* P5 | 5’-TTTCGGAACTGCCTTGGTGGCGG-3’ |
| *atu3779* P6 | 5’-ATTCCCGGCCGGAACTACCATCGC-3’ |
| *atu2112* P1 | 5’- AAAAAGCTTTATTTCGTCTTCGAGGATGGG-3’ |
| *atu2112* P2 | 5’- GATGATCGATATGACAGAAACAGTGAAATGGC-3’ |
| *atu2112* P3 | 5’-ATCGATCATCTGCGAAATTGCG-3’ |
| *atu2112* P4 | 5’-AAAGAATTCATATCGTCCTCGGTTTCCGC-3’ |
| *atu2112* P5 | 5’- AAAAAGCTTTATTTCGTCTTCGAGGATGGG-3’ |
| *atu2112* P6 | 5’- TTTGCACCGAAAGATGCCGCG-3’ |
| *atu1022* P1 | 5’-AAAAAGCTTTTATGCGCTTTGACCAGCGCACCC-3’ |

| | |
|---|---|
| *atu1022* P2 | 5’-AATCCCCAGACTGTCTTTTTCATGCCG-3’ |
| *atu1022* P3 | 5’-TGGGGATTATTCAGGCACGGGCTAGCC-3’ |
| *atu1022* P4 | 5’-AAAGAATTCAATACGCTCTTCAACTCCATCCG-3’ |
| *atu1022* P5 | 5’- AAAAGCGACGTCGCTCGCG-3’ |
| *atu1022* P6 | 5’-AAACTACTACGACGAAGACGGTCAGG-3’ |
| *atu2117* P1 | 5’-AAAGAATTCAAGAGAATGTCTGGACAGGCGTGGC-3’ |
| *atu2117* P2 | 5’-ATTTTCAATCTAGAGTTGCCGCCTTCGTTATGCC-3’ |
| *atu2117* P3 | 5’- TTGAAAATGCTGCGGCTCCC-3’ |
| *atu2117* P4 | 5’-AAAAAGCTTTAGATGTTCCTGTCGAACACCG-3’ |
| *atu2117* P5 | 5’-ATACAGGCGCATCGCGGCC-3’ |
| *atu2117* P6 | 5’-AAGGCGAGGTGGTCACTGACC -3’ |
| *atu3093* P1 | 5’-GCTGCAACTAGTGCGCCACAGCCCATCTCG-3’ |
| *atu3093* P2 | 5’-AAGCATGGTACCGAATTCGCGGCTTGTTGATATTCCT-3’ |
| *atu3093* P3 | 5’-GAATTCGGTACCATGCTTTAGCAGCGGCGGGCGCCGGA-3’ |
| *atu3093* P4 | 5’-GCACGTAAGCTTGCGGTGATCTTGATGAT-3’ |
| *atu3093* P5 | 5’-GAGCTTCTCGGGCAATTCCG-3’ |
| *atu3093* P6 | 5’-GCCTCACGAAGCCGCACGATC-3’ |

| | |
|---|---|
| *atu3077* NdeI Fwd | 5’- GCGGCGCATATGAAATTTAATCGCAGACAT-3' |
| *atu3077 NdeI BamHI Rvs* | 5’- GTCGTCGGATCCTTATTGCGCCTTTTCGAG-3' |
| *atu3078* NdeI Fwd | 5’- GCGGCGCATATGGTTCGGCAATTTCTTCCC-3' |
| *atu3078 BamHI* Rvs | 5’- GTCGTCGGATCCTCACGCCGTTCCGATGGT-3' |
| *atu3779 NdeI* Fwd | 5’- GCGGCGCATATGACAAAGACCCTTTCAAAT-3' |
| *atu3779 KpnI* Rvs | 5’- GCGGCGCATATGACAAAGACCCTTTCAAAT-3' |
| *atu2113 NdeI* Fwd | 5’- GCGGCGCATATGAAAGAATGTCTGCCGGAT-3' |
| *atu2113 BamHI Rvs* | 5’- GTCGTCGGATCCTCAGGCGGCCGAAAATCT-3' |

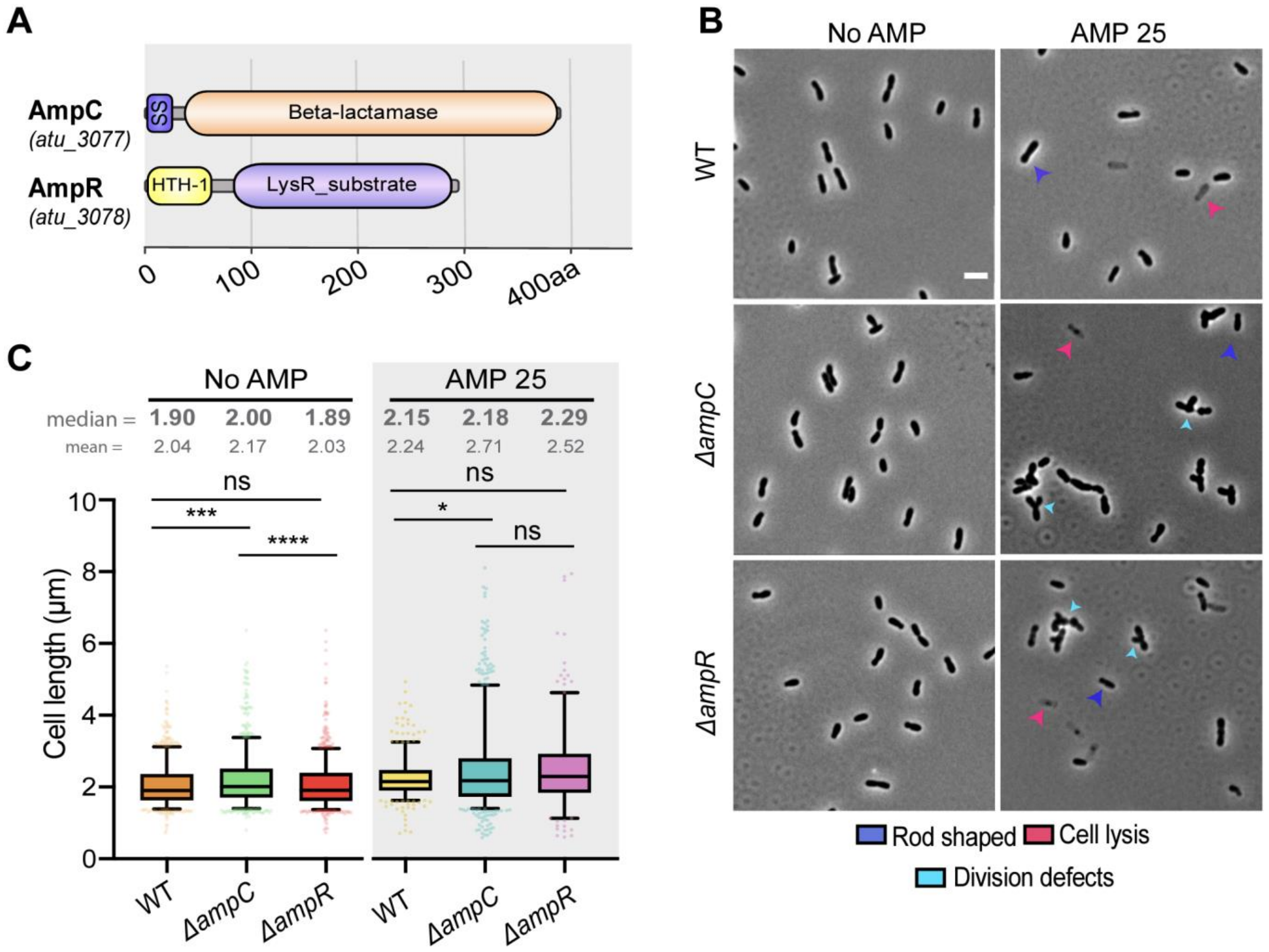

**Supplemental 1**

**SUPPLEMENTAL FIG 1. Putative enzymes involved in ampicillin resistance in the plant pathogen, *A. tumefaciens* C58. (A)** Schematic representation of putative proteins involved in ampicillin resistance. Corresponding ORF numbers are indicated under the name of the protein. **(B)** Phase-contrast microscopy of exponentially growing WT *A. tumefaciens* C58 and *ΔampC* and *ΔampR* incubated in the absence (No AMP) or presence of ampicillin 25 µg/ml (AMP 25) for 2 h. Scale bar = 2 µm. Arrows indicate major phenotypes; Blue = WT/Rod-shaped, magenta = cell lysis, cyan = division defects. **(C)** Box-plots represent the distribution of cell length in WT *A. tumefaciens* C58, *ΔampC,* and *ΔampR* of untreated cells (No AMP) and treated with ampicillin 25 µg/ml (AMP 25) for 2 h before imaging. Top number represents the median cell length. Top quartile (Q1) indicates 75% of the population, middle line represents the median, and bottom quartile (Q2) represents the 25% of the population. Whiskers indicate 5-95% of population. Significance of the cell length distributions are indicated using the non-parametric Kruskal-Wallis test followed by a Dunn posthoc analysis (* = $P<0.1$; *** = $P<0.001$; **** = $P<0.0001$; ns = not significant).

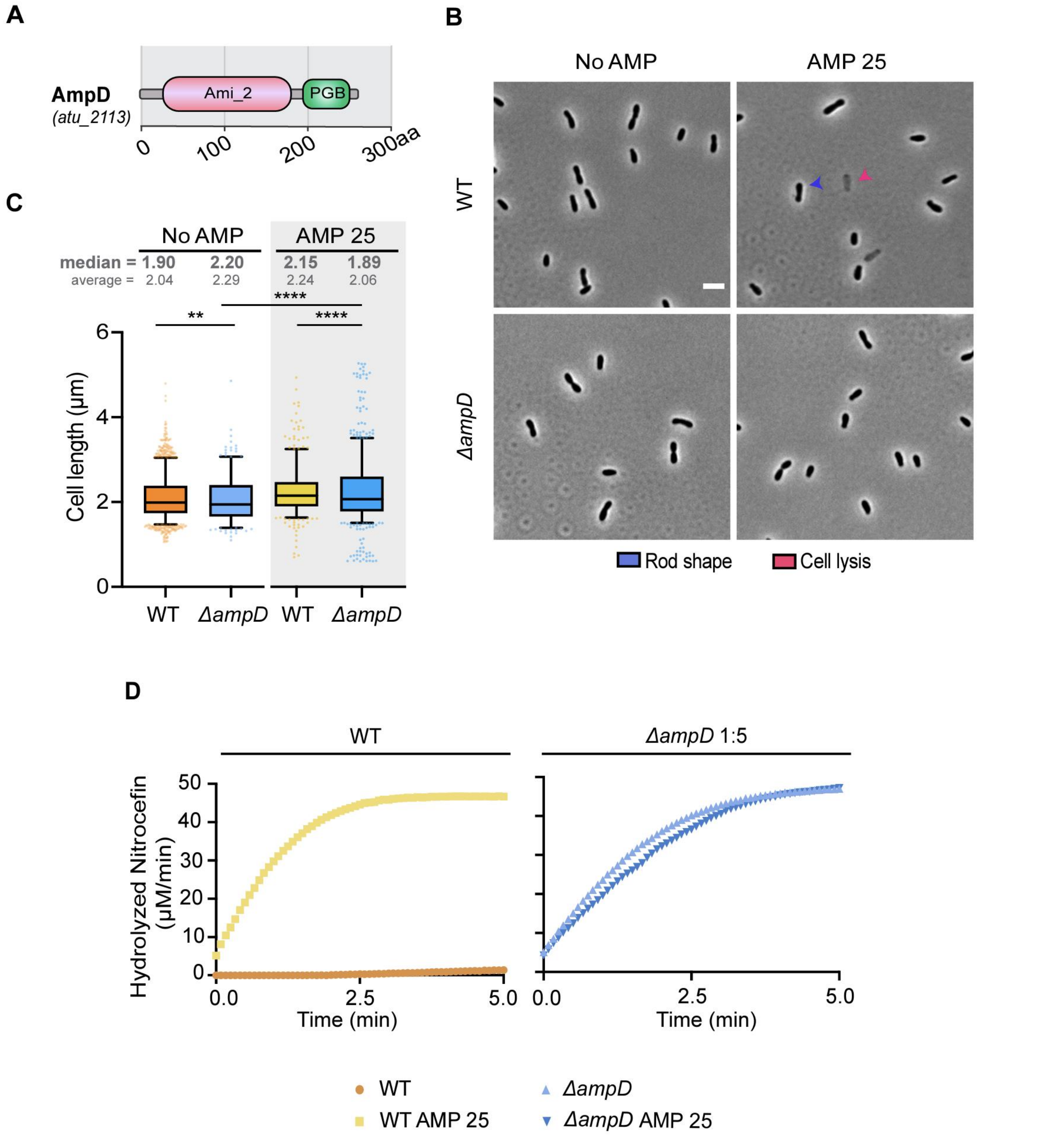

**Supplemental 2**

**SUPPLEMENTAL FIG 2. Loss of AmpD leads to ampicillin resistance. (A)** Schematic representation of putative proteins involved in ampicillin resistance. Corresponding ORF numbers are indicated under the name of the protein. **(B)** Phase-contrast microscopy of exponentially growing WT *A. tumefaciens* C58 and *ΔampD* incubated in the absence (No AMP) or presence of ampicillin 25 µg/ml (AMP 25) for 2 h. Scale bar = 2 µm. Arrows indicate major phenotypes; Blue = WT/Rod-shaped, magenta = cell lysis. **(C)** Box-plots represent the distribution of cell length in WT *A. tumefaciens* C58 and *ΔampD* of untreated cells (No AMP) and treated with ampicillin 25 µg/ml (AMP 25) for 2 h before imaging. Distribution of cell lengths was obtained using MicrobJ (87). Top number represents the median cell length. Top quartile (Q1) indicates 75% of the population, middle line represents the median, and bottom quartile (Q2) represents the 25% of the population. Whiskers indicate 5-95% of population. Significance of the cell length distributions are indicated using the non-parametric Kruskal-Wallis test followed by a Dunn posthoc analysis (** = $P < 0.1$; ** = $P < 0.01$, *** = $P < 0.001$, **** = $P < 0.0001$*). **(D)** Determination of β-lactamase production was performed via a nitrocefin assay using cell lysates. “No AMP” or “AMP 25” indicates cells untreated or treated, respectively, with ampicillin 25 µg/ml for 2 h before the generation of cell lysates. *ΔampD* lysates were normalized based on total protein content (7.5 µg/ml) and subsequently diluted to 1:5 as the rate of nitrocefin hydrolysis was significantly faster than the controls (WT, WT AMP 25).

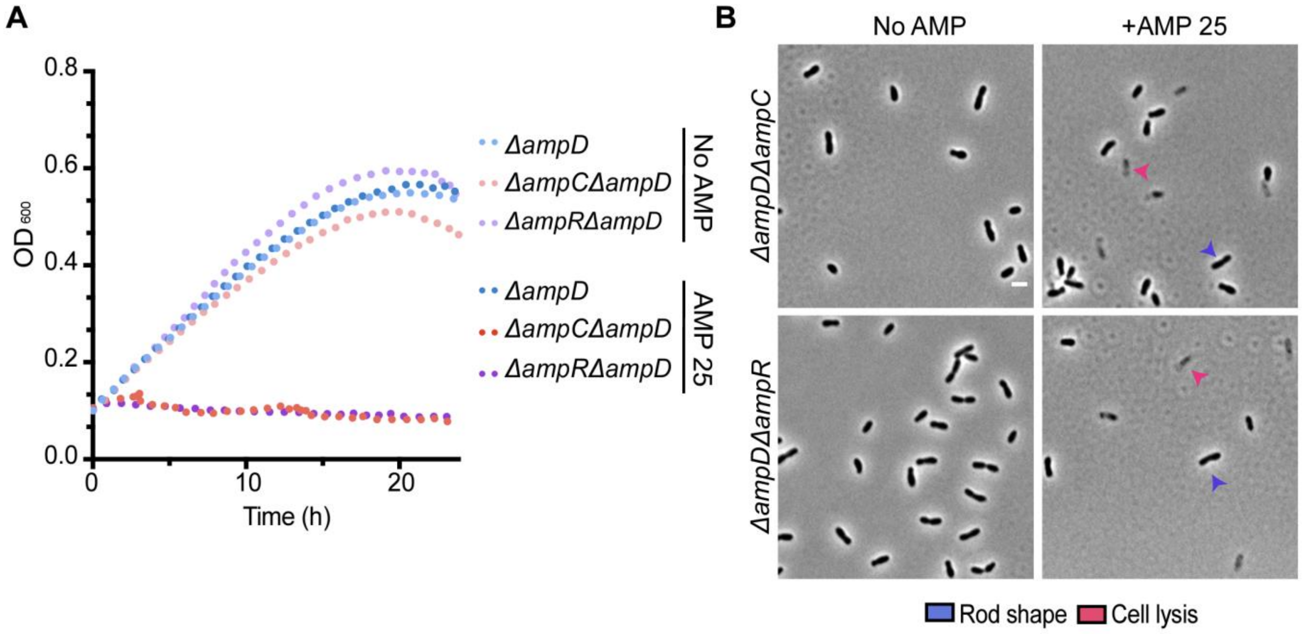

**Supplemental 3**

**SUPPLEMENTAL FIG 3. Loss of AmpD results in constitutive β-lactamase production activity and elevated ampicillin resistance. (A)** Growth of *A. tumefaciens* strains in the absence (No AMP) and presence (AMP) of ampicillin 25 µg/ml (AMP 25) for 24 h (n=1, 2 replicates). (**B)** Phase-contrast microscopy of exponentially growing strains treated with ampicillin 25 µg/ml (AMP 25) for 2 h.

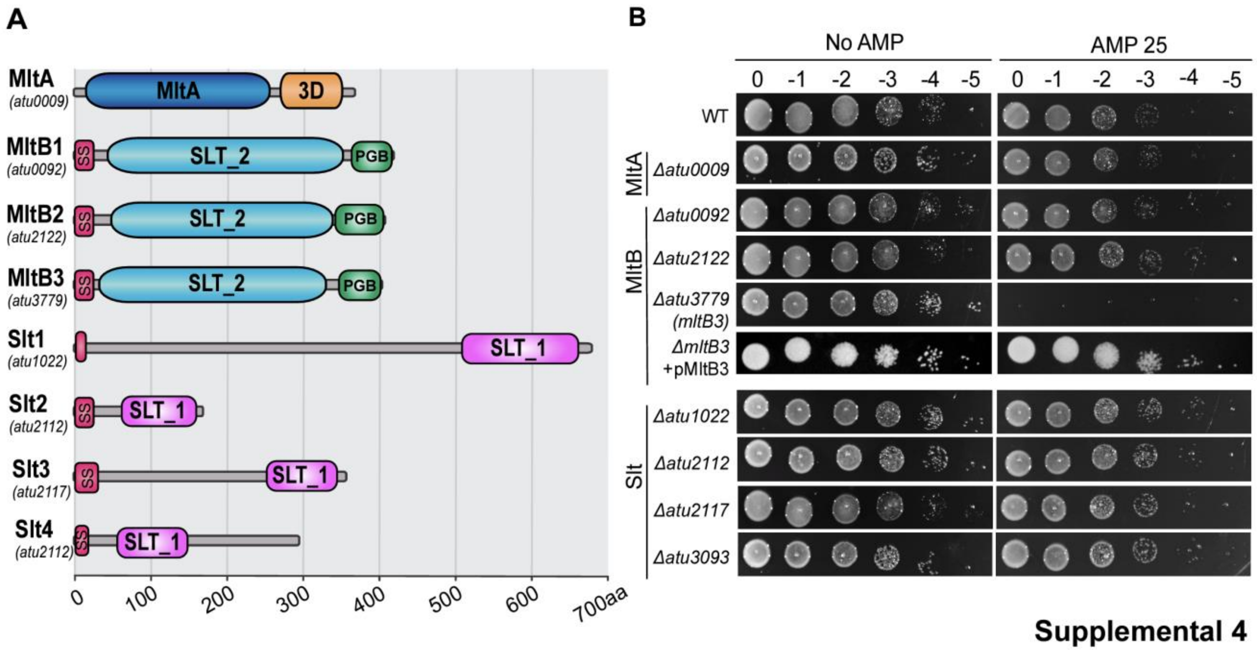

**Supplemental 4**

**SUPPLEMENTAL FIG 4. MltB3 is required for *A. tumefaciens* natural resistance to ampicillin. (A)** Domain organization of predicted lytic transglycosylase (LT) proteins. **(B)** Ampicillin susceptibility assay was performed via spotting dilutions. Briefly, indicated strains were cultured ON at 28°C, serially diluted, spotted on solid medium containing no ampicillin (No AMP) or ampicillin 25 µg/ml (AMP 25), and incubated at 28C for 36 h before imaging. Plates used to demonstrate complementation of *ΔmltB3* (*ΔmltB3* +pMltB3) included 1µM IPTG to induce expression of plasmid encoded MltB**.**
